# Diversity and gene expression patterns of functional groups in sidestream and mainstream wastewater partial-nitritation anammox biofilms

**DOI:** 10.1101/2020.04.23.057489

**Authors:** Carolina Suarez, David Gustavsson, Malte Hermansson, Frank Persson

## Abstract

Partial nitritation-anammox (PNA) is today used for nitrogen removal from highly concentrated wastewater after anaerobic sludge digestion (sidestream). However, implementation of PNA for treatment of municipal wastewater (mainstream), with its lower ammonium concentration and lower temperature is challenging, which might be due to differences in microbial community composition and/or activity. To investigate this, we compared side-by-side sidestream and mainstream PNA biofilms using amplicon sequencing of 16S rDNA and rRNA, *hzsB* DNA and mRNA, and the genes *nxrB*, and *amoA*. The two communities were different to each other with relatively more heterotrophic denitrifying bacteria and less anammox bacteria in the mainstream. With *hzsB* and *nxrB* we found microdiversity among *Brocadia* and *Nitrospira*, and turnover (taxa replacement) between sidestream and mainstream. However, in both environments *Brocadia sapporoensis* represented most of the *hzsB* DNA and mRNA reads, despite the different environmental conditions and nitrogen removal rates. All of those populations present in both sidestream and mainstream had no differences in their 16S rRNA:rDNA ratios, supporting recent findings that rRNA:rDNA ratios are poor indicators of bacterial activity. The observed diversity within functional groups and composition differences between sidestream and mainstream add complexity to our view of PNA communities with possible implication for reactor function.

## Introduction

Excess of reactive nitrogen in the environment contributes to eutrophication (Erisman, *et al.* 2015). To reduce reactive nitrogen discharges into water bodies, removal of nitrogen in wastewater treatment plants (WWTPs) is essential. Biological nitrogen removal from the sidestream of wastewater, i.e. reject water, from dewatering of anaerobic digested sludge, with high ammonium concentration and high temperature, can be achieved by the partial nitritation-anammox (PNA) process (Lackner, *et al*. 2014). PNA combines oxidation of part of the wastewater ammonium to nitrite by ammonia oxidising bacteria (AOB) and a subsequent conversion of the nitrite and remaining ammonium to nitrogen gas by anammox bacteria (AMX). PNA communities are often grown in biofilms in granule reactors or in moving bed biofilm reactors (MBBRs) to maintain the slow growing AMX at high concentrations in the reactors (Agrawal, *et al.* 2017).

Implementation of PNA for the colder, more diluted mainstream of wastewater, which contains the majority of the nitrogen at WWTPs, has been challenging. Low nitrogen removal rates and high NO_3_^-^ production are commonly reported (Gonzalez-Martinez, *et al.* 2016, Gustavsson, *et al.* 2020, Lotti, *et al.* 2014, Wu, *et al.* 2016). From a population ecology perspective, multiple scenarios exist to explain the differences in removal rates and nitrate production between sidestream and mainstream PNA. First, the lower substrate concentration and the lower temperature in the mainstream will inevitably result in lower removal rates. Second, some taxa could differ in their metabolic activity in the two environments. For example, high nitrate production by nitrite oxidising bacteria (NOB) is often reported in mainstream PNA (Cao, *et al.* 2017). Third, functional groups might differ in abundance, which in turn could influence ecosystem function. When gradually replacing sidestream with mainstream wastewater a decrease in AMX and AOB abundances was observed by Yang, *et al.* (2018). Fourth, diversity in the accessory genome within bacterial species exist (McInerney, *et al.* 2017). A mechanism explaining microdiversity is that sub-populations have different ecological niches, i.e. are ecotypes (Moore, *et al.* 1998). It is possible that differences within the main functional groups in sidestream and mainstream PNA could occur. For instance, cold tolerant strains with an oligotrophic lifestyle might be observed in mainstream, while sidestream conditions might favour eutrophic lifestyles.

Amplicon sequencing of the 16S gene (rDNA) has recently been employed to investigate the effect of various operational conditions in PNA systems, and to describe community composition (Agrawal, *et al*. 2017, Laureni, *et al.* 2016, Persson, *et al.* 2017, Yang, *et al.* 2018). Sequencing of rDNA offers limited resolution to infer closely related populations, but the use of amplicon sequences variants (ASVs) (Callahan, *et al.* 2017) instead of operational taxonomic units (OTUs) would allow potential ecotypes to be elucidated (García-García, *et al.* 2019). Even higher resolution within taxonomic groups could be achieved by sequencing of functional genes, like *hzsB* for anammox bacteria (Wang, *et al.* 2012), *amoA* for AOB (Rotthauwe, *et al.* 1997), and *nxrB* for NOB (Pester, *et al.* 2013), but this approach is rarely used for describing the PNA communities.

Community composition can be described with amplicon sequencing, but bacteria can be active, growing, dormant or deceased (Blazewicz, *et al.* 2013), and their metabolic status cannot be determined from gene sequencing alone. An alternative is sequencing of 16S rRNA (rRNA), as bacterial growth has been associated with an increase in ribosome production, at least for *Proteobacteria* (Kerkhof and Kemp 1999, Schaechter, *et al.* 1958), and ribosomal degradation is seen for some bacteria during starvation (Deutscher 2003). Thus, estimations of rRNA:rDNA ratios have been considered a measure of activity in the total community (Campbell, *et al.* 2011, Jones and Lennon 2010). However, this metric is not universal, as there are exceptions to the link between rRNA content and activity in bacteria (Blazewicz, *et al.* 2013).

In this study, we operated pilot-scale MBBRs for PNA, fed with either pre-treated municipal wastewater (mainstream) or sludge liquor from anaerobic sludge digesters (sidestream) from the Sjölunda WWTP, Malmö, Sweden. The aim of this study was to determine, by high throughput amplicon sequencing of rDNA, if different microbial communities and/or taxa abundance were established in the mainstream and sidestream. We also asked if multiple AMX, NOB and AOB populations coexisted in the two environments. To address this question, we sequenced the functional genes *hzsB, nxrB* and *amoA*, as well as *hzsB* mRNA from the biofilm on individual mainstream and sidestream carriers. Furthermore, we measured rRNA:rDNA ratios to tests our hypothesis that the rRNA:rDNA ratios of bacterial groups present in both environments would vary due to the different conditions, which would provide insights about the response in activity of specific taxa.

## Materials and Methods

The sidestream and mainstream pilot MBBRs were located at the Sjölunda WWTP, Malmö, Sweden (Hanner, *et al.* 2003), see Table 1 for operational data. They were filled to 40% with K1^®^ carriers (Veolia Water Technologies AB – AnoxKaldnes, Lund, Sweden). The MBBRs are described in detail elsewhere (Gustavsson, *et al.* 2020). To promote anammox growth in the mainstream MBBR, biofilm carriers were frequently exchanged between the sidestream and mainstream MBBRs. However, biofilm carriers sampled in this study were not exchanged, but kept isolated in each reactor in cylindrical cages (immersed volume 2.5 L) for 128 days until sampling. A steel mesh bottom in the cages allowed water circulation. The cages were filled using carries taken from their respective MBBRs.

**Table 1:** Operational data of the pilot reactors from 15 September 2015 to 14 October 2015. Mean values +/- S.D.

|  | Sidestream | Mainstream |
| --- | --- | --- |
| Influent $\text{NH}_4^+$ (mg N $\text{L}^{-1}$) | $880 \pm 48$ | $26 \pm 5.0$ |
| Effluent $\text{NH}_4^+$ (mg N $\text{L}^{-1}$) | $55 \pm 12$ | $21 \pm 4.6$ |
| Effluent $\text{NO}_2^-$ (mg N $\text{L}^{-1}$) | $6.1 \pm 2.1$ | $0.25 \pm 0.52$ |
| Effluent $\text{NO}_3^-$ (mg N $\text{L}^{-1}$) | $120 \pm 13$ | $1.7 \pm 1.3$ |
| Ammonium loading rate (g N $\text{m}^{-2} \text{d}^{-1}$) | $1.8 \pm 0.25$ | $0.78 \pm 0.14$ |
| Nitrogen removal rate (g N $\text{m}^{-2} \text{d}^{-1}$) | $1.5 \pm 0.19$ | $0.13 \pm 0.12$ |
| DO (mg $\text{L}^{-1}$) | $1.0 \pm 0.24$ | $1.0 \pm 0.15$ |
| T ($^{\circ}\text{C}$) | $28.3 \pm 0.80$ | $19.1 \pm 0.65$ |

### Sampling

Biofilm carriers from each cage were snap-frozen in an ethanol-dry ice mixture immediately at sampling, kept frozen in dry ice during transportation and then stored at −80°C. Sidestream biofilms had a red colour, while a brown colour was observed in the mainstream biofilms (Figure S1, Supporting information). In addition, a wet weight of 431 mg ± 36 (average ± 95% confidence interval) was observed for sidestream biofilms, and 296 ± 13 for mainstream biofilms.

### Co-extraction of DNA and RNA

Carriers with biofilms were thawed in RNA later-ICE (Thermo Fisher Scientific, Waltham, MA USA). The biofilm was removed from the carrier compartments and added to a lysis matrix tube E (MP biomedicals, Santa Ana, CA, USA) with 800 μl of lysis solution of a ZR-duet MiniPrep kit (Zymo Research). Mechanical disruption of the biofilm was done with a FastPrep-24 5G (MP biomedicals) at speed 6 for 40 seconds. Subsequent steps of the DNA-RNA co-extraction were carried out with the ZR-duet kit according to manufacturer instructions. Ribolock RNase inhibitor (40 U/μl; Thermo Fisher Scientific) was added to the extracted RNA. DNA and RNA concentration were measured after extraction using a Qubit 3.0 fluorometer (Thermo Fisher Scientific). Potential traces of genomic DNA were removed from the RNA extraction with a DNA-free DNA Removal kit (Thermo Fisher Scientific). cDNA was synthesised with SuperScript VILO MasterMix (Thermo Fisher Scientific) according to the manufacturer’s instructions.

### Sequencing

PCR amplification of the 16S V4 region was done with primers 515’F (Hugerth, *et al.* 2014) and 806R (Caporaso, *et al.* 2011), using dual indexing of the primers (Kozich, *et al.* 2013). Template DNA (40 ng) or undiluted cDNA (2 μl) was amplified in a total volume of 50 μl using a Phusion Hot Start II DNA Polymerase (Thermo Fisher Scientific). The following PCR program was used: activation (98°C, 30 s); 30 cycles of denaturation (98°C, 10 s), annealing (56°C, 30 s) and elongation (72°C, 15 s); followed by final elongation (72°C, 10 min). PCR products were purified with Ampure XP (Beckman Coulter, Brea, CA, USA). Purified PCR products were pooled in equimolar amounts. Sequencing was performed on an Illumina MiSeq using the MiSeq Reagent Kit v3 (Illumina, San Diego, CA, USA).

PCR amplification of *nxrB*, *hzsB*, and *amoA* was carried out in two steps using Nextera index adapters (Illumina). Primers nxrB169F and nxrB638R (Pester, *et al.* 2013) were used for *nxrB* amplification, amplification of *hzsB* was done with the hzsB_396F and hzsB_742R primers (Wang, *et al.* 2012), and primers AmoA1F mod (Stephen, *et al.* 1999) and AmoA2R (Rotthauwe, *et al.* 1997) were used for *amoA*. The following PCR program was used for all amplicons: activation (98°C, 30 s); 25 cycles of denaturation (98°C, 10 s), annealing (56°C, 30 s) and elongation (72°C, 45 s); followed by final elongation (72°C, 5 min). PCR amplicons were then purified, and a second PCR with 8 cycles of amplification was used to attach the index adapters followed by a second purification. Amplicons were then pooled together in equimolar amounts and sequenced on a MiSeq as described above.

### Statistics and data analysis

Samples with less than 30,000 reads were excluded prior to analysis. ASVs were generated, using DADA2 version 1.12 (Callahan, *et al.* 2016). The SILVA 132 database (Quast, *et al.* 2013) was used for taxonomic classification of the 16S amplicons with IDTAXA (Murali, *et al.* 2018). Data was analysed in R (R Core Team 2019) using the packages Phyloseq (McMurdie and Holmes 2013) and Vegan (Oksanen, *et al*. 2019). Raw sequence reads were deposited at the NCBI (Bioproject: PRJNA552732). See table S1 for individual accession numbers of each sample.

Reads were normalized by proportion prior to estimation of beta diversity between mainstream and sidestream in the rDNA, rRNA, *nxrB* and *hzsB* libraries and for comparisons between the rDNA and rRNA libraries. Beta diversity was estimated with the abundance-based Bray-Curtis index and the presence-absence-based Simpson index, which is not sensitive to richness differences, and is a measure of turnover (Baselga 2010). To estimate differential abundance of ASVs between sidestream and mainstream, DESeq2 was used (Love, *et al*. 2014) without subsampling before the analysis (McMurdie and Holmes 2014). A p_(adj)_ <0.01 value (DESeq2) was used as criterion for statistical significance. Only samples with both rDNA and rRNA libraries available (mainstream, n=6; sidestream, n=7) were used for plots of rDNA vs rRNA. Ratios of rRNA:rDNA were estimated after excluding values of zero in the libraries.

### Fluorescence in situ hybridization (FISH)

FISH was carried out on suspended biomass as previously described (Suarez, *et al.* 2015). The probe AMX820 (Schmid, *et al.* 2001), labelled with FAM, was used to targ *et al* l *Brocadia* populations. The probes BAN162 (Schmid, *et al.* 2001), labelled with Cy3, and BFU613 (van de Vossenberg, *et al.* 2008), labelled with Cy5, were used to target *Brocadia* subpopulations, as they only partially cover the *Brocadia* genus, as determined with ARB 6.0.6 (Ludwig, *et al.* 2004) using the SILVA NR 132 SSU *Brocadia* sequences. These probes were applied together with unlabelled competitors (for Ban162, CGG TAG CCC CAA TTG CTT; for Bfu613, GGA TGC CGT TCT TCC GTT GAG CGG) to increase probe specificity, as previously reported (Persson, *et al.* 2014, Suarez, *et al.* 2015).

## Results and disscusion

### Sidestream and mainstream communities

We compared PNA microbial communities exposed to sidestream and mainstream conditions by sequencing rDNA and rRNA from individual biofilm carriers. Mainstream and sidestream communities, estimated from both rDNA and rRNA, were significantly different, as shown with the abundance-based Bray-Curtis index (Adonis β_bray_; p < 0.01, r^2^ = 0.16; Figure 1A). Because beta-diversity can also exist among rare taxa (Gobet, *et al*. 2012), we also used the presence-absence Simpson index, which measures species turnover (Baselga 2010). The observed results (Adonis β_Sim_; p <0.01, r^2^ = 0.32; Figure 1B), suggest that the sidestream and mainstream communities not only differed significantly in their relative abundance of taxa, but also in their identity. The sidestream and mainstream communities also diverged from the initial seed communities, indicating temporal dynamics (Figure S2, Supporting information). Although elucidating the assembly mechanism of PNA biofilms is beyond the scope of this study, sidestream and mainstream were exposed to different environmental conditions and subjected to potential immigration from two different water sources, which would both influence the community composition.

**Figure 1:**
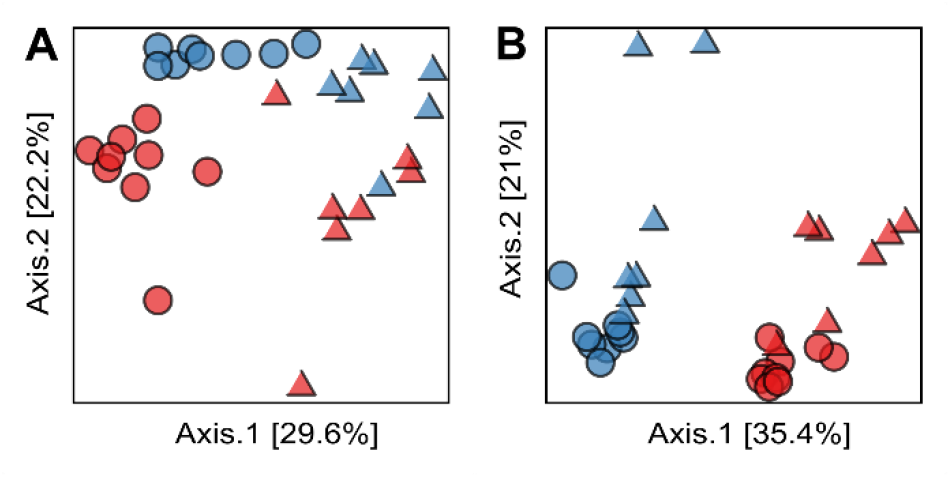
PCoA of rDNA and rRNA libraries, based on the abundance-based Bray-Curtis index (**A**) and the presence-absence-based Simpson index (**B**). Red: mainstream, Blue: sidestream. Circles: rDNA, triangles: rRNA.

### Comparing rDNA and rRNA libraries

The rDNA and rRNA libraries of the mainstream and sidestream communities were different in relative read abundance (Adonis β_bray_; p < 0.001, r ^2^= 0.24; Figure 1A), which could be interpreted as rRNA:rDNA ratios different to one. However, turnover between the rDNA and rRNA libraries was also observed (Adonis β_sim_; p < 0.001, r^2^ = 0.13; Figure 1B), which would imply that different ASVs were detected by sequencing of rDNA and rRNA. ASVs present in the rRNA, but not the rDNA libraries are known as phantom taxa. They could be the result of PCR errors during reverse transcription, but could also arise due to rDNA under-sampling of rare but highly active taxa (Klein, *et al*. 2016). Supporting the latter suggestion, we commonly observed phantom taxa for genera with high rRNA:rDNA values, like *Competibacter*, *Agitococcus, Romboutsia*, the AMX *Brocadia* and the AOB *Nitrosomonas* (Figure S3, Supporting information). In addition, among genera with low rRNA:rDNA, like *Denitratisoma, Dokdonella* and *UTBCD1* (Figure S3, Supporting information), some ASVs did not have any corresponding rRNA reads. This could signal that a high proportion of these taxa in the biofilms were dormant or dead. Extracellular DNA is commonly observed in biofilms (Dominiak, *et al*. 2011) and could inflate rDNA reads for some taxa (Albertsen, *et al*. 2015).

Despite the differences, relative read abundances of the rDNA and rRNA libraries over the entire dataset were positively correlated at the phylum level (Kendall’s τ = 0.77, p < 0.001, Figure 2) as well as the ASV level (Kendall’s τ = 0.59, p < 0.001), and also seen by modelling rRNA abundance of ASVs with beta regression (z =18.0, p<0.001, pseudo-R^2^ = 0.44; Figure S4, Supporting information). Thus, ribosomal relative abundance was generally linked to taxa relative abundance. Nonetheless, it appears that rRNA:rDNA ratios varied between phyla; for example, low ratios were observed for *Acidobacteria*, and high ratios were noticed for *Firmicutes* and *Planctomycetes* (Figure 2, S4, table S2). Such different rRNA:rDNA ratios have been observed before across phyla (Denef, *et al.* 2016, Steven, *et al.* 2017).

**Figure 2.**
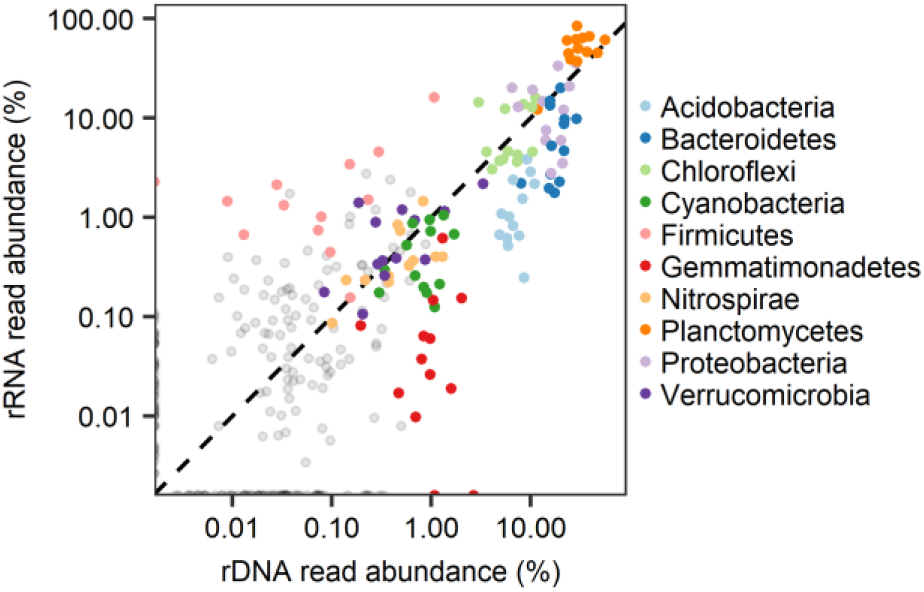
Comparison of rDNA and rRNA of the entire dataset at the phylum-level. Each point indicates the abundance of an ASV in one MBBR carrier; the black dashed diagonal line indicates equal rDNA and rRNA abundance. The colours denote the 10 most abundant phyla.

*Acidobacteria* are considered slow growing bacteria (Ward, *et al*. 2009), and hence it is possible that their low rRNA:rDNA ratios in fact represent slow growth rates compared to other taxa in the biofilms. This might be supported by the fact that *Acidobacteria* have between one and two copies of the 16S RNA gene, since low copy numbers are associated with oligotrophic lifestyles (Klappenbach, *et al.* 2000, Stevenson and Schmidt 2004). On the other hand, although rRNA:rDNA values larger than one are frequently used as criteria for activity (Blazewicz, *et al.* 2013), different taxa might differ in their rRNA content during growth or dormancy to the extent that rRNA content and growth rate are not linked (Blazewicz, *et al.* 2013). Thus, potentially active taxa might be misclassified as dormant just because their rRNA:rDNA ratio is below one (Steven, *et al.* 2017).

The disproportionally high read abundances of rRNA at low rDNA abundances for most phyla (Figure S4) confirm previous similar observations, suggesting higher activity among rare taxa (Campbell, *et al.* 2011, Jia, *et al.* 2019, Jones and Lennon 2010, Klein, *et al.* 2016, Wilhelm, *et al.* 2014). This phenomenon could perhaps be due to under-sampling (Steven, *et al.* 2017), but higher growth rate among rare taxa could arise due to intraspecific competition or predation of abundant taxa (Jousset, *et al*. 2017). In fact, by using metagenomic data, Jia, *et al*. (2019) observed higher replication rates for taxa at low relative abundances supporting that this is a real phenomenon.

### Comparing rRNA:rDNA ratios between sidestream and mainstream

We originally expected that the same ASVs present in the relatively different sidestream and mainstream conditions would have different rRNA:rDNA ratios. For example, in the case of *Brocadia* we anticipated a higher rRNA:rDNA in the sidestream because the activity in terms of nitrogen conversion was much higher than in mainstream (Table 1). In addition, the growth rate of anammox bacteria is affected by temperature (Laureni, *et al.* 2015). But differences in rRNA:rDNA ratios between the two environments were not observed for any of the 12 ASVs present across all rRNA and rDNA libraries (Wilcoxon rank sum test, p>0.05; Figure 3).

**Figure 3:**
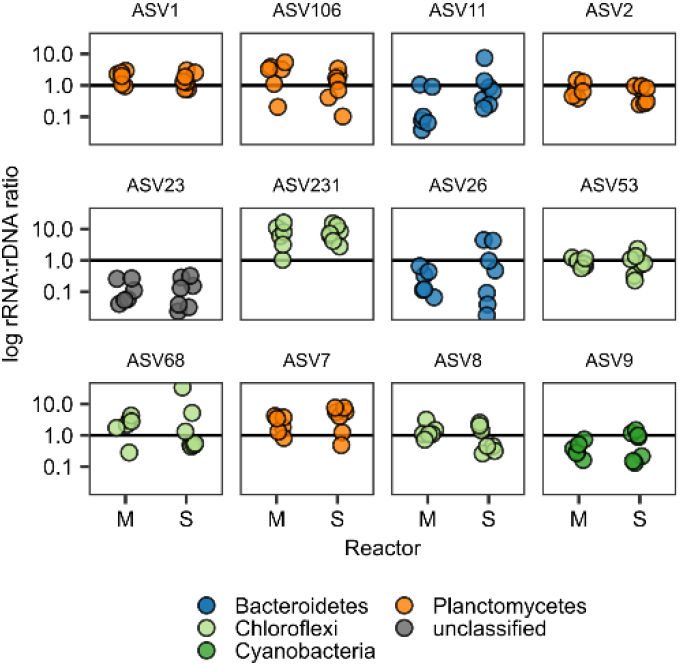
Ratios of rRNA:rDNA for 12 ASVs present in all rDNA and rRNA libraries. Colours denote phylum classification, M = mainstream and S = sidestream. ASV1 and ASV7 are classified as the AMX *Brocadia*.

We conclude that, at least for PNA systems, the rRNA:rDNA ratios are not a direct proxy for metabolic activity. This would agree with recent studies in soil using stable isotope probing showing that rRNA:rDNA ratios underestimate active taxa and are a poor predictor of rRNA synthesis (Papp, *et al.* 2018a, Papp, *et al.* 2018b). For some taxa, ribosomes may not be degraded during slow/no growth conditions. For *Nitrosomonas*, stable ribosomal content has been observed during inhibition (Wagner, *et al.* 1995) and preservation of old ribosomes appears to occur in *Thaumarchaeota* (Papp, *et al.* 2019). Furthermore, presence of rRNA is not necessarily an indication of active rRNA production, therefore it might be useful to complement measurements of rRNA:rDNA ratios with other methods such as stable isotope probing (Papp, *et al.* 2018a).

### Sidestream and mainstream communities

As also reported for other PNA MBBRs (Agrawal, *et al*. 2017, Persson, *et al*. 2017), the dominant taxa in the biofilms were AMX (Figure 4A), with *Brocadia* being the only AMX genus detected. Relative read abundance of *Brocadia* rDNA was higher in the sidestream than the mainstream biofilms (DESeq2; P_(adj)_< 0.01; Figure 4B). Furthermore, potential heterotrophic denitrifying bacteria(HDB) like *Zoogloea* and *Sulfuritalea*, among others, were more abundant in the Mainstream (DESeq2; p_(adj)_ < 0.01, Figure 4B) and were in fact more or less absent in the sidestream biofilms. Similarly, Yang, *et al.* (2018) observed a decrease in AMX abundance when a sidestream community was exposed to mainstream conditions, also indicating that sidestream conditions favour AMX growth relative to other bacteria. The larger fraction of HDB in the mainstream suggests more extensive competition for NO_2_^-^ between AMX and HDB; in addition, potential nitrogen loops of nitrification, anammox, denitrification and DNRA may occur (Speth, *et al.* 2016).

**Figure 4.**
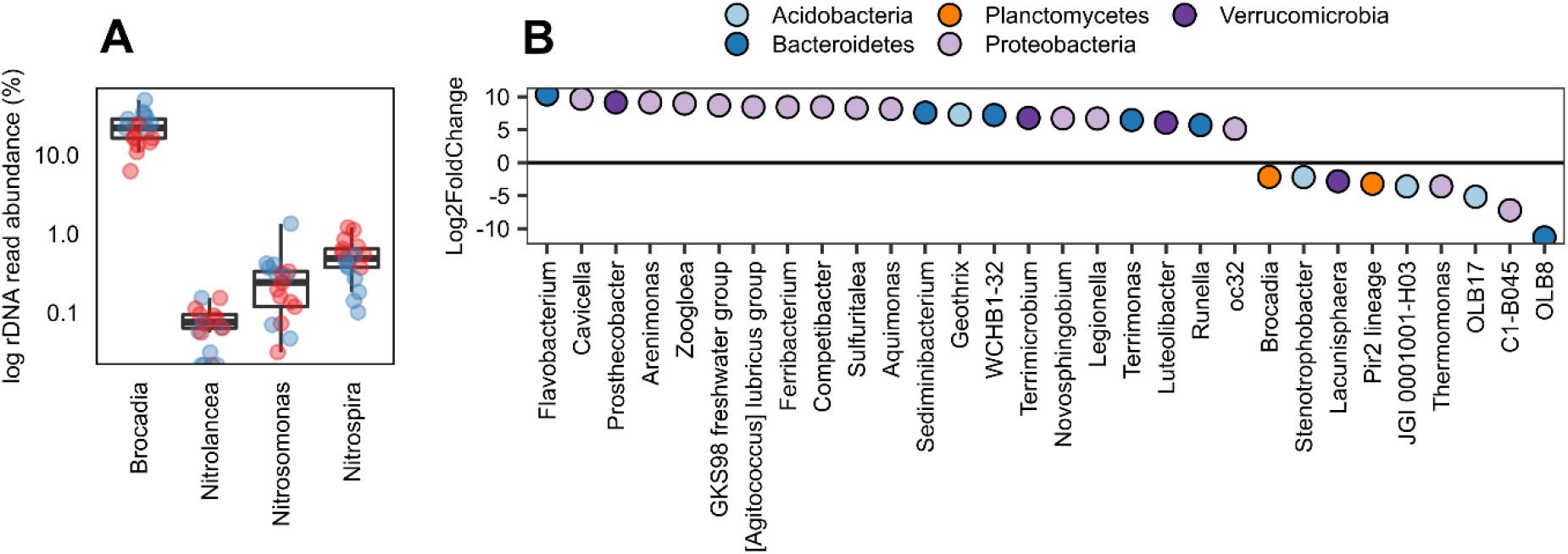
Abundance of taxa at the genus level. **A:** rDNA read abundance of AMX (*Brocadia*), AOB (*Nitrosomonas*) and NOB (*Nitrospira and Nitrolancea*), red: mainstream, blue: sidestream. **B:** Log2FoldChange (LFC) for rDNA genera with differential abundance between mainstream and sidestream biofilms (DESeq2; p_(adj)_ < 0.01). Positive LFC corresponds to higher abundance in mainstream, and negative LFC corresponds to higher abundance in sidestream; only the top 30 genera with the lowest significant p(adj) are shown.

### Microdiversity of nitrogen transforming bacteria

Several anammox populations coexisted in the biofilms as assessed by rDNA sequencing (Figure S5-S7, Supporting information). An example of the microdiversity within *Brocadia* can be observed in Figure 5A, where a combination of different FISH probes was used to visualize three different *Brocadia* subpopulations. By sequencing the *hzsB* gene, 119 ASVs within *Brocadia* were detected (Figure 5B, S8, Supporting information), and turnover between sidestream and mainstream was demonstrated (Adonis, β_sim_, r^2^=0.57, p=0.01). Nonetheless, a single ASV was dominant in both sidestream and mainstream (Figure 5B, S8, Supporting information), and represented 77 ± 3% of the total *hzsB* reads (Figure 5C). As hydrazine synthase (HZS) is a key enzyme for the anammox process, this suggests that this strain was responsible for the bulk anaerobic oxidation of ammonium in both the sidestream and the mainstream, in spite of the different environmental conditions in the reactors.

**Figure 5A:**
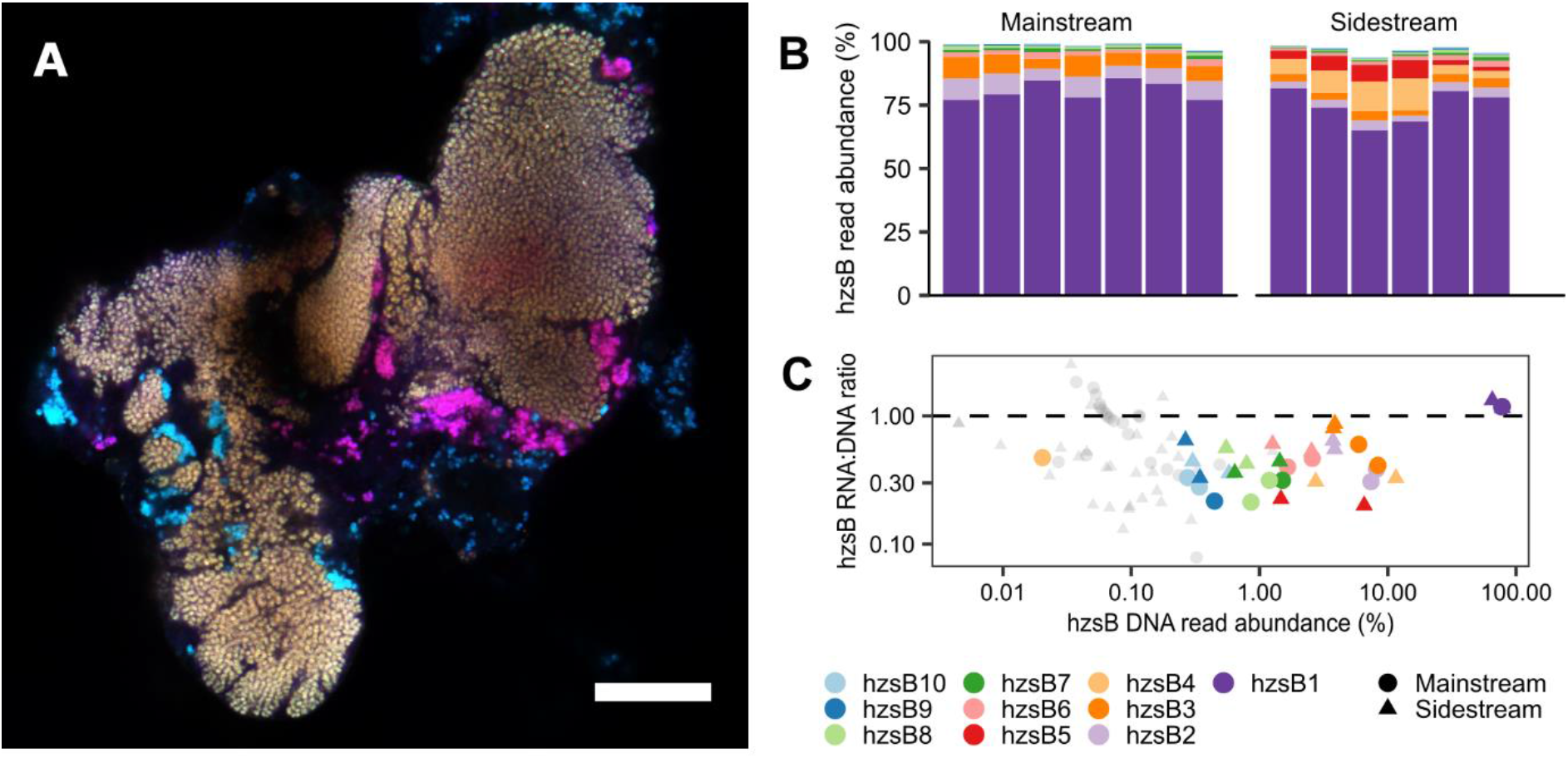
Multiple *Brocadia* populations (sidestream) targeted with the FISH probes AMX820 (Blue), BFU613 (Red) and Ban162 (Green); overlap among probes results in additional colours: Cyan (AMX820 and Ban162), Magenta: (AMX820, BFU613), White: (AMX820, Ban162 and BFU613). **B**: DNA read abundance of the top 10 *hzsB* ASVs; each colour represents a unique ASV; each bar represents a biofilm sample. **C:** RNA:DNA ratios for the top 10 *hzsB* ASVs (n=4). Same colour coding as in B.

Using rDNA, several *Nitrospira* and *Nitrosomonas* ASVs were observed (Figure S6, S7, Supporting information). Likewise, using the key enzymes *nxrB* and *amoA* we also observed multiple *Nitrospira* ASVs (Figure 6, S9, Supporting information) and *Nitrosomonas* ASVs (Figure S10, Supporting information), respectively. Turnover of *Nitrospira* communities in sidestream and mainstream was observed as assessed with *nxrB* (Adonis, β_sim_, p=0.004, r^2^=0.39), while for *amoA*, low PCR yield in sidestream samples prevented comparison of sidestream and mainstream. In microbial communities the coexistence of closely related taxa is often reported (Goldford, *et al.* 2018). For example, for *Nitrospira*, which is commonly present in wastewater (Daims, *et al.* 2001), multiple populations can coexist in activated sludge (Gruber-Dorninger, *et al.* 2014). Coexistence of multiple AMX populations has also been reported in PNA systems (Bhattacharjee, *et al*. 2017, Laureni, *et al*. 2019, Persson, *et al*. 2014). *Nitrospira* is a heterogeneous group, which not only represents a nitrite oxidising potential, but also contains populations capable of using a myriad of electron donors and - acceptors (Daims, *et al.* 2015, Koch, *et al.* 2015, van Kessel, *et al.* 2015) and the same has also been shown for AMX (Hu, *et al*. 2019, Kartal, *et al*. 2007, Strous, *et al*. 2006, van de Vossenberg, *et al*. 2008) and AOB (Bock, *et al.* 1995, Schmidt, *et al.* 2004). Thus, the coexistence of different *Brocadia, Nitrosomonas* and *Nitrospira* ASVs, observed in this study, could be a general phenomenon, explained by ecotypes within the AMX, AOB and the NOB utilizing different metabolic pathways and playing various ecological roles. The roles of the less abundant AMX in the PNA biofilms are yet unclear, but read abundance of *hzsB* mRNA (Figure 5C) suggests that their contribution to the anammox process might have been minor.

**Figure 6:**
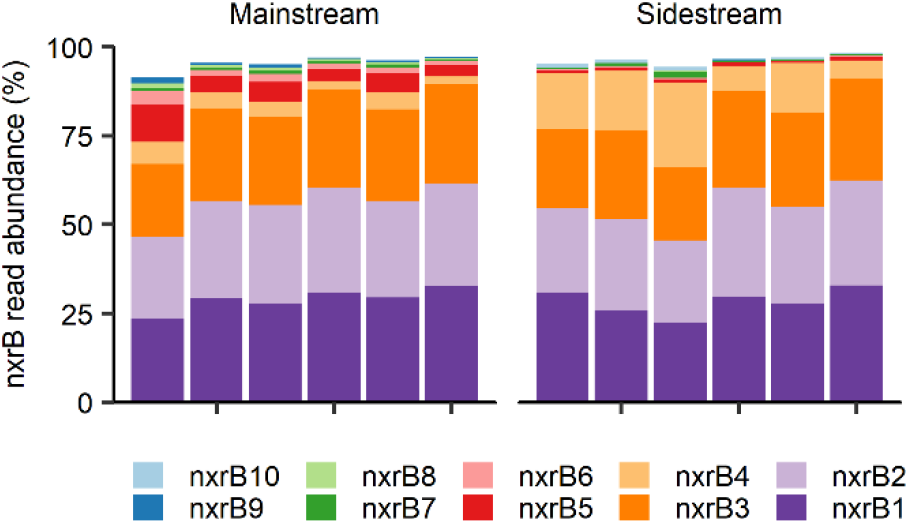
Read abundance of the top 10 *nxrB* ASVs.

The dominant *hzsB* ASV had an identical sequence to *B. sapporoensis*, which is frequently observed in PNA bioreactors (Lotti, *et al.* 2015b, Persson, *et al.* 2014) and is considered relatively fast growing (Lotti, *et al.* 2015a). It has a maximum specific anammox activity at 37°C (Narita, *et al.* 2017), which, together with other factors like nitrogen load and partial nitritation rate, would explain the lower nitrogen removal rates observed in the mainstream compared with the sidestream MBBR. A considerable decrease of anammox rates with decreasing temperature has often been described in PNA reactors (Gilbert, *et al.* 2014, Laureni, *et al.* 2016, Lotti, *et al.* 2015b). Interestingly, no particular cold-tolerated AMX have so far been detected, instead AMX within *Brocadia* are commonly reported in mainstream reactors. This study suggests that *B. sapporoensis*, can in fact be competitive relative to other AMX taxa at both sidestream and mainstream conditions. One possible explanation for such a phenomenon can be mechanisms of adaptation, where temperature changes can result in changes in membrane composition (Rattray, *et al*. 2010) and protein expression (Lin, *et al*. 2018). Alternatively, there could be multiple closely related populations with identical *hzsB* and rDNA sequences, but with different accessory genes, and thus metagenomic studies would be needed to resolve such populations.

Comammox can occur among lineage II in *Nitrospira* (Daims, *et al.* 2015). The majority of *Nitrospira* in this study were from lineage I (Figure S9, Supporting information) and thus the comammox process was likely not important for the nitrogen transformations in the biofilm. Inhibition of NOB is critical for the function of PNA, and several strategies have been proposed for NOB suppression including, but not limited to, DO limitation, intermittent aeration and exposure to free nitrous acid (Malovanyy, *et al.* 2015, Pérez, *et al.* 2014, Wang, *et al.* 2016). However, the observed NOB diversity within *Nitrospira* could impact reactor performance, because microbial diversity might lead to functional redundancy (Allison and Martiny 2008). As NOB in any PNA reactor likely consist of multiple coexisting populations, operational strategies to reduce nitrite oxidation based on knowledge gained from pure cultures or enrichments may not be adequate. In fact, the inhibition of a dominant NOB population, may well lead to the succession of another dominant population. Future studies, combining well-defined reactor experiments with high resolution genomics would gain further insights in how microdiversity of key guilds, as observed here, affects turnover of substrates and thereby reactor performance

## Funding

This work was supported by FORMAS [245-2014-1528, 942-2015-683 and 2018-01423], Wilhelm och Martina Lundgrens vetenskapsfond and Adlerbertska forskningsstiftelsen.

## Acknowledgements

The authors acknowledge the Genomics core facility at the University of Gothenburg, the Centre for Cellular Imaging at the University of Gothenburg and the National Microscopy Infrastructure, NMI (VR-RFI 2016-00968), for providing support and use of their equipment, and the colleagues at the Sjölunda WWTP, for monitoring the pilot reactors.

## Notes

### Competing Interest Statement

The authors have declared no competing interest.

## References

Agrawal S, Karst Søren M, Gilbert Eva M et al. The role of inoculum and reactor configuration for microbial community composition and dynamics in mainstream partial nitritation anammox reactors. MicrobiologyOpen 2017;6: e00456.

Albertsen M, Karst SM, Ziegler AS et al. Back to Basics – The Influence of DNA Extraction and Primer Choice on Phylogenetic Analysis of Activated Sludge Communities. PLoS ONE 2015;10: e0132783.

Allison SD, Martiny JBH. Resistance, resilience, and redundancy in microbial communities. Proc Natl Acad Sci USA 2008;105: 11512–9.

Baselga A. Partitioning the turnover and nestedness components of beta diversity. Glob Ecol Biogeogr 2010;19: 134–43.

Bhattacharjee AS, Wu S, Lawson CE et al. Whole-Community Metagenomics in Two Different Anammox Configurations: Process Performance and Community Structure. Environ Sci Technol 2017;51: 4317–27.

Blazewicz SJ, Barnard RL, Daly RA et al. Evaluating rRNA as an indicator of microbial activity in environmental communities: limitations and uses. ISME J 2013;7: 2061–8.

Bock E, Schmidt I, Stüven R et al. Nitrogen loss caused by denitrifying Nitrosomonas cells using ammonium or hydrogen as electron donors and nitrite as electron acceptor. Arch Microbiol 1995;163: 16–20.

Callahan BJ, McMurdie PJ, Holmes SP. Exact sequence variants should replace operational taxonomic units in marker-gene data analysis. ISME J 2017;11: 2639–43.

Callahan BJ, McMurdie PJ, Rosen MJ et al. DADA2: High-resolution sample inference from Illumina amplicon data. Nat Meth 2016;13: 581–3.

Campbell BJ, Yu L, Heidelberg JF et al. Activity of abundant and rare bacteria in a coastal ocean. Proc Natl Acad Sci USA 2011;108: 12776–81.

Cao Y, van Loosdrecht MCM, Daigger GT. Mainstream partial nitritation-anammox in municipal wastewater treatment: status, bottlenecks, and further studies. Appl Microbiol Biotechnol 2017;101: 1365–83.

Caporaso JG, Lauber CL, Walters WA et al. Global patterns of 16S rRNA diversity at a depth of millions of sequences per sample. Proc Natl Acad Sci USA 2011;108: 4516–22.

Daims H, Lebedeva EV, Pjevac P et al. Complete nitrification by *Nitrospira* bacteria. Nature 2015;528: 504–9.

Daims H, Nielsen JL, Nielsen PH et al. In Situ Characterization of Nitrospira-Like Nitrite-Oxidizing Bacteria Active in Wastewater Treatment Plants. Appl Environ Microbiol 2001;67: 5273–84.

Denef VJ, Fujimoto M, Berry MA et al. Seasonal Succession Leads to Habitat-Dependent Differentiation in Ribosomal RNA:DNA Ratios among Freshwater Lake Bacteria. Front Microbiol 2016;7: 606.

Deutscher MP. Degradation of Stable RNA in Bacteria. J Biol Chem 2003;278: 45041–4.

Dominiak DM, Nielsen JL, Nielsen PH. Extracellular DNA is abundant and important for microcolony strength in mixed microbial biofilms. Environ Microbiol 2011;13: 710–21.

Erisman JW, Galloway JN, Dise NB et al. Nitrogen: too much of a vital resource: Science Brief. WWF science brief NL. Zeist, The Netherlands: WWF Netherlands, 2015.

García-García N, Tamames J, Linz AM et al. Microdiversity ensures the maintenance of functional microbial communities under changing environmental conditions. ISME J 2019;13: 2969–83.

Gilbert EM, Agrawal S, Karst SM et al. Low Temperature Partial Nitritation/Anammox in a Moving Bed Biofilm Reactor Treating Low Strength Wastewater. Environ Sci Technol 2014;48: 8784–92.

Gobet A, Böer SI, Huse SM et al. Diversity and dynamics of rare and of resident bacterial populations in coastal sands. ISME J 2012;6: 542–53.

Goldford JE, Lu N, Bajić D et al. Emergent simplicity in microbial community assembly. Science 2018;361: 469–74.

Gonzalez-Martinez A, Rodriguez-Sanchez A, Garcia-Ruiz MJ et al. Performance and bacterial community dynamics of a CANON bioreactor acclimated from high to low operational temperatures. Chem Eng J 2016;287: 557–67.

Gruber-Dorninger C, Pester M, Kitzinger K et al. Functionally relevant diversity of closely related Nitrospira in activated sludge. ISME J 2014;9: 643–55.

Gustavsson DJI, Suarez C, Wilén B-M et al. Long-term stability of partial nitritation-anammox for treatment of municipal wastewater in a moving bed biofilm reactor pilot system. Sci Total Environ 2020;714: 136342.

Hanner N, Aspegren H, Nyberg U et al. Upgrading the Sjölunda WWTP according to a novel process concept. Water Sci Technol 2003;47: 1–7.

Hu Z, Wessels HJCT, van Alen T et al. Nitric oxide-dependent anaerobic ammonium oxidation. Nature Comm 2019;10: 1244.

Hugerth LW, Wefer HA, Lundin S et al. DegePrime, a Program for Degenerate Primer Design for Broad-Taxonomic-Range PCR in Microbial Ecology Studies. Appl Environ Microbiol 2014;80: 5116–23.

Jia Y, Leung MHY, Tong X et al. Rare Taxa Exhibit Disproportionate Cell-Level Metabolic Activity in Enriched Anaerobic Digestion Microbial Communities. mSystems 2019;4: e00208–18.

Jones SE, Lennon JT. Dormancy contributes to the maintenance of microbial diversity. Proc Natl Acad Sci USA 2010;107: 5881–6.

Jousset A, Bienhold C, Chatzinotas A et al. Where less may be more: how the rare biosphere pulls ecosystems strings. ISME J 2017;11: 853–62.

Kartal B, Kuypers MMM, Lavik G et al. Anammox bacteria disguised as denitrifiers: nitrate reduction to dinitrogen gas via nitrite and ammonium. Environ Microbiol 2007;9: 635–42.

Kerkhof L, Kemp P. Small ribosomal RNA content in marine Proteobacteria during non-steady-state growth. FEMS Microbiol Ecol 1999;30: 253–60.

Klappenbach JA, Dunbar JM, Schmidt TM. rRNA Operon Copy Number Reflects Ecological Strategies of Bacteria. Appl Environ Microbiol 2000;66: 1328–33.

Klein AM, Bohannan BJM, Jaffe DA et al. Molecular Evidence for Metabolically Active Bacteria in the Atmosphere. Front Microbiol 2016;7: 772.

Koch H, Lücker S, Albertsen M et al. Expanded metabolic versatility of ubiquitous nitrite-oxidizing bacteria from the genus *Nitrospira*. Proc Nat Acad Sci 2015;112: 11371–6.

Kozich JJ, Westcott SL, Baxter NT et al. Development of a Dual-Index Sequencing Strategy and Curation Pipeline for Analyzing Amplicon Sequence Data on the MiSeq Illumina Sequencing Platform. Appl Environ Microbiol 2013;79: 5112–20.

Lackner S, Gilbert EM, Vlaeminck SE et al. Full-scale partial nitritation/anammox experiences-An application survey. Water Res 2014;55: 292–303.

Laureni M, Falås P, Robin O et al. Mainstream partial nitritation and anammox: long-term process stability and effluent quality at low temperatures. Water Res 2016;101: 628–39.

Laureni M, Weissbrodt DG, Szivák I et al. Activity and growth of anammox biomass on aerobically pre-treated municipal wastewater. Water Res 2015;80: 325–36.

Laureni M, Weissbrodt DG, Villez K et al. Biomass segregation between biofilm and flocs improves the control of nitrite-oxidizing bacteria in mainstream partial nitritation and anammox processes. Water Res 2019;154: 104–16.

Lin X, Wang Y, Ma X et al. Evidence of differential adaptation to decreased temperature by anammox bacteria. Environ Microbiol 2018;20: 3514–28.

Lotti T, Kleerebezem R, Abelleira-Pereira JM et al. Faster through training: The anammox case. Water Res 2015a;81: 261–8.

Lotti T, Kleerebezem R, Hu Z et al. Pilot-scale evaluation of anammox-based mainstream nitrogen removal from municipal wastewater. Environ Technol 2015b;36: 1167–77.

Lotti T, Kleerebezem R, Hu Z et al. Simultaneous partial nitritation and anammox at low temperature with granular sludge. Water Res 2014;66: 111–21.

Love MI, Huber W, Anders S. Moderated estimation of fold change and dispersion for RNA-seq data with DESeq2. Genome Biol 2014;15: 550.

Ludwig W, Strunk O, Westram R et al. ARB: a software environment for sequence data. Nucleic Acids Res 2004;32: 1363–71.

Malovanyy A, Yang J, Trela J et al. Combination of upflow anaerobic sludge blanket (UASB) reactor and partial nitritation/anammox moving bed biofilm reactor (MBBR) for municipal wastewater treatment. Bioresour Technol 2015;180: 144–53.

McInerney JO, McNally A, O’Connell MJ. Why prokaryotes have pangenomes. Nature Microbiol 2017;2: 17040.

McMurdie PJ, Holmes S. phyloseq: An R Package for Reproducible Interactive Analysis and Graphics of Microbiome Census Data. PLoS ONE 2013;8: e61217.

McMurdie PJ, Holmes S. Waste Not, Want Not: Why Rarefying Microbiome Data Is Inadmissible. PLoS Comput Biol 2014;10: e1003531.

Moore LR, Rocap G, Chisholm SW. Physiology and molecular phylogeny of coexisting Prochlorococcus ecotypes. Nature 1998;393: 464–7.

Murali A, Bhargava A, Wright ES. IDTAXA: a novel approach for accurate taxonomic classification of microbiome sequences. Microbiome 2018;6: 140.

Narita Y, Zhang L, Kimura Z-i et al. Enrichment and physiological characterization of an anaerobic ammonium-oxidizing bacterium ‘Candidates Brocadia sapporoensis’. Syst Appl Microbiol 2017;40: 448–57.

Oksanen J, Blanchet FG, Friendly M et al. vegan: Community Ecology Package, 2019.

Papp K, Hungate BA, Schwartz E. Microbial rRNA Synthesis and Growth Compared through Quantitative Stable Isotope Probing with H218O. Appl Environ Microbiol 2018a;84: e02441–17.

Papp K, Hungate BA, Schwartz E. mRNA, rRNA and DNA quantitative stable isotope probing with H218O indicates use of old rRNA among soil Thaumarchaeota. Soil Biol Biochem 2019;130: 159–66.

Papp K, Mau RL, Hayer M et al. Quantitative stable isotope probing with H218O reveals that most bacterial taxa in soil synthesize new ribosomal RNA. ISME J 2018b;12: 3043–5.

Pérez J, Lotti T, Kleerebezem R et al. Outcompeting nitrite-oxidizing bacteria in single-stage nitrogen removal in sewage treatment plants: A model-based study. Water Res 2014;66: 208–18.

Persson F, Suarez M, Hermansson M et al. Community structure of partial nitritation-anammox biofilms at decreasing substrate concentrations and low temperature Microb Biotechnol 2017;10: 761–72.

Persson F, Sultana R, Suarez M et al. Structure and composition of biofilm communities in a moving bed biofilm reactor for nitritation-anammox at low temperatures. Bioresour Technol 2014;154: 267–73.

Pester M, Maixner F, Berry D et al. NxrB encoding the beta subunit of nitrite oxidoreductase as functional and phylogenetic marker for nitrite-oxidizing Nitrospira. Environ Microbiol 2013;16: 3055–71.

Quast C, Pruesse E, Yilmaz P et al. The SILVA ribosomal RNA gene database project: improved data processing and web-based tools. Nucleic Acids Res 2013;41: D590–D6.

R Core Team. R: A Language and Environment for Statistical Computing. Vienna, Austria: R Foundation for Statistical Computing, 2019.

Rattray JE, van de Vossenberg J, Jaeschke A et al. Impact of Temperature on Ladderane Lipid Distribution in Anammox Bacteria. Appl Environ Microbiol 2010;76: 1596–1603.

Rotthauwe JH, Witzel KP, Liesack W. The ammonia monooxygenase structural gene amoA as a functional marker: molecular fine-scale analysis of natural ammonia-oxidizing populations. Appl Environ Microbiol 1997;63: 4704–12.

Schaechter M, MaalØe O, Kjeldgaard NO. Dependency on Medium and Temperature of Cell Size and Chemical Composition during Balanced Growth of Salmonella typhimurium. Microbiol 1958;19: 592–606.

Schmid M, Schmitz-Esser S, Jetten M et al. 16S-23S rDNA intergenic spacer and 23S rDNA of anaerobic ammonium-oxidizing bacteria: implications for phylogeny and in situ detection. Environ Microbiol 2001;3: 450–9.

Schmidt I, van Spanning RJM, Jetten MSM. Denitrification and ammonia oxidation by Nitrosomonas europaea wild-type, and NirK- and NorB-deficient mutants. Microbiol 2004;150: 4107–14.

Speth DR, In ‘t Zandt MH, Guerrero-Cruz S et al. Genome-based microbial ecology of anammox granules in a full-scale wastewater treatment system. Nat Comm 2016;7: 11172.

Stephen JR, Chang Y-J, Macnaughton SJ et al. Effect of Toxic Metals on Indigenous Soil β-Subgroup Proteobacterium Ammonia Oxidizer Community Structure and Protection against Toxicity by Inoculated Metal-Resistant Bacteria. Appl Environ Microbiol 1999;65: 95–101.

Steven B, Hesse C, Soghigian J et al. Simulated rRNA/DNA Ratios Show Potential To Misclassify Active Populations as Dormant. Appl Environ Microbiol 2017;83: e00696–17.

Stevenson BS, Schmidt TM. Life History Implications of rRNA Gene Copy Number in Escherichia coli. Appl Environ Microbiol 2004;70: 6670–7.

Strous M, Pelletier E, Mangenot S et al. Deciphering the evolution and metabolism of an anammox bacterium from a community genome. Nature 2006;440: 790–4.

Suarez C, Persson F, Hermansson M. Predation of nitritation-anammox biofilms used for nitrogen removal from wastewater. FEMS Microbiol Ecol 2015;91: fiv124.

van de Vossenberg JLCM, van Niftrik L, Strous M et al. Candidatus ‘Brocadia fulgida’: an autofluorescent anaerobic ammonium oxidizing bacterium. FEMS Microbiol Ecol 2008;63: 46–55.

van Kessel MAHJ, Speth DR, Albertsen M et al. Complete nitrification by a single microorganism. Nature 2015;528: 555–9.

Wagner M, Rath G, Amann R et al. In situ Identification of Ammonia-oxidizing Bacteria. Syst Appl Microbiol 1995;18: 251–64.

Wang D, Wang Q, Laloo A et al. Achieving Stable Nitritation for Mainstream Deammonification by Combining Free Nitrous Acid-Based Sludge Treatment and Oxygen Limitation. Sci Rep 2016;6: 25547.

Wang Y, Zhu G, Harhangi HR et al. Co-occurrence and distribution of nitrite-dependent anaerobic ammonium and methane-oxidizing bacteria in a paddy soil. FEMS Microbiol Lett 2012;336: 79–88.

Ward NL, Challacombe JF, Janssen PH et al. Three genomes from the phylum Acidobacteria provide insight into the lifestyles of these microorganisms in soils. Appl Environ Microbiol 2009;75: 2046–56.

Wilhelm L, Besemer K, Fasching C et al. Rare but active taxa contribute to community dynamics of benthic biofilms in glacier-fed streams. Environ Microbiol 2014;**16**: 2514–24.

Wu S, Bhattacharjee AS, Weissbrodt DG et al. Effect of short term external perturbations on bacterial ecology and activities in a partial nitritation and anammox reactor. Bioresour Technol 2016;219: 527–35.

Yang Y, Zhang L, Cheng J et al. Microbial community evolution in partial nitritation/anammox process: From sidestream to mainstream. Bioresour Technol 2018;251: 327–33.

