## Supporting information for "Diversity and gene expression patterns of functional groups in sidestream and mainstream wastewater partial-nitritation anammox biofilms"

### Suplementary methods

For a subset of the 16S dataset of abundant phyla, the relationship between rDNA and rRNA was analysed with a beta regression model (Ferrari & Cribari-Neto, 2004) using the R package betareg (Cribari-Neto & Zeileis, 2010) and a logit link function. In addition to rDNA as independent variable, we included phylum and their interaction with rDNA as categorical variables, with *Proteobacteria* as the reference level for Phylum. We used both rDNA and phylum as a covariate for the precision parameter ($\phi$).

To estimate phylogeny of *amoA*, *hzsB* and *nxrB*, references were obtained by blastx against the nr NCBI database among *Nitrosomandales*, *Brocadiales* and *Nitrospirae* respectively; uncultured/environmental sample sequences were excluded. The reading frame of nucleotide sequences was corrected with DECIPHER (Wright, 2016) prior to translation into amino acid sequences with the R package Biostrings (Pagès *et al.*, 2019). *Nitrospira amoA*, *Scalindua hzsB* and anammox *nxrB* were used as outgroups. Amino acid alignments were generated with DECIPHER. Maximum likelihood trees were estimated using a WAG+G4 substitution model with the Phangorn package (Schliep, 2010). A few ASVs corresponding to proteins with premature stop codons were observed for *nxrB* and *hzsB* and were excluded prior to analysis. Bootstrap support was assessed by 500 re-samplings. Phylogenetic trees were visualized with the Interactive tree of life web tool (Letunic & Bork, 2019).

### Supplementary figures

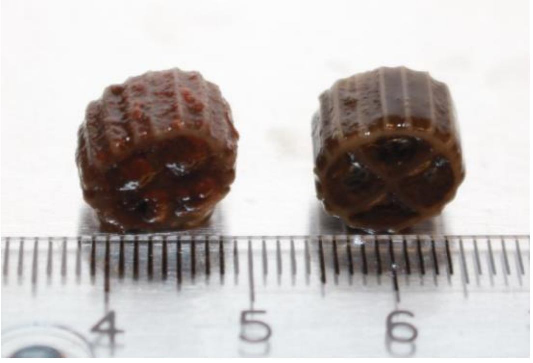


**Figure S1:** A: Biofilm carriers from sidestream (left) and mainstream (right); a ruler in cm is shown for size comparison.

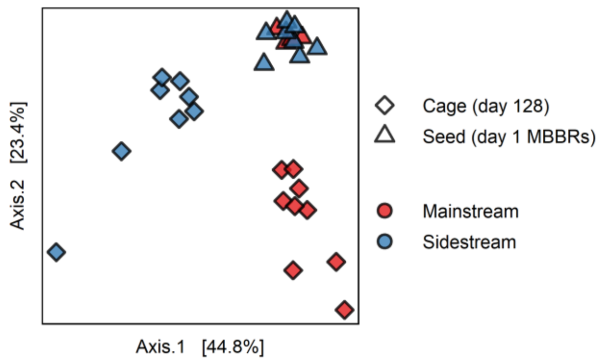


**Figure S2:** PCoA of rDNA libraries, based the Simpson index for the seed (day 1) and cage (day128) samples. Seed are the mature biofilms taken directly from the MBBRs.

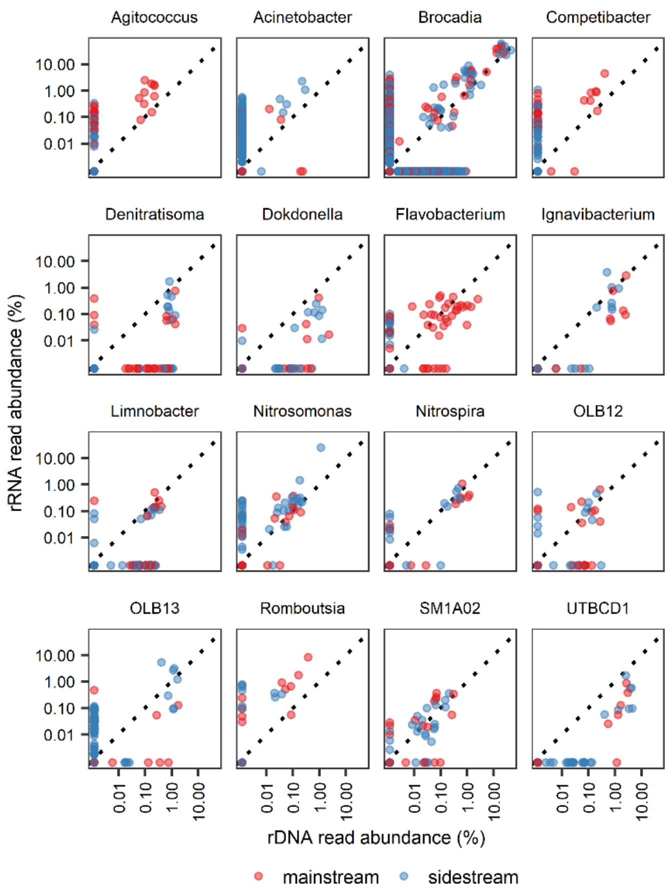


**Figure S3:** Comparison of rDNA and rRNA at the ASV level for selected genera. A point indicates abundance of an ASV in one MBBR carrier; the diagonal line indicates equal rDNA and rRNA abundance. Red: Mainstream. Blue: Sidestream. A small value (0.001) was added to both rDNA and rRNA to plot values of zero in a log scale.

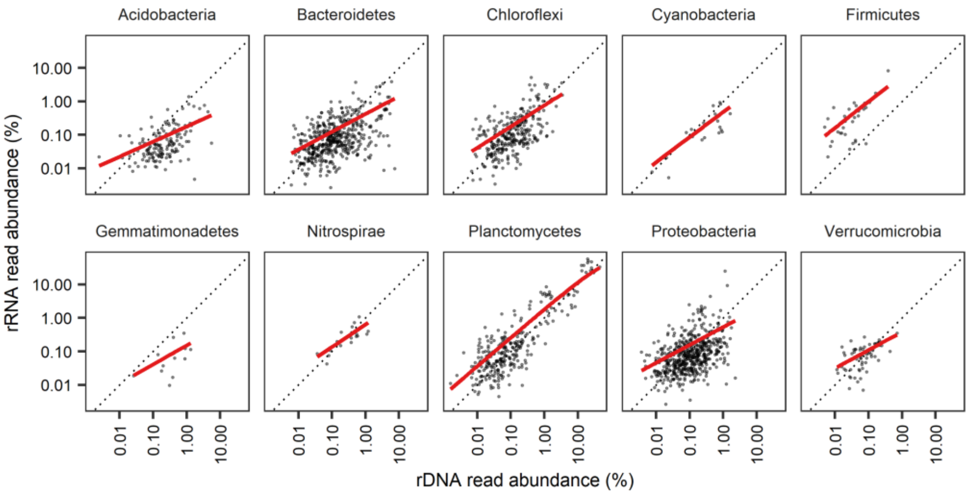


**Figure S4.** Comparison of rDNA and rRNA of at the ASV level grouped by Phylum. A point indicates abundance of an ASV in one MBBR carrier. The thick lines are the mean values predicted by a beta regression (see Table S2 for statistics). The black dashed diagonal line indicates equal rDNA and rRNA abundance (rRNA:rDNA value of one). Values of zero rDNA and rRNA are excluded.

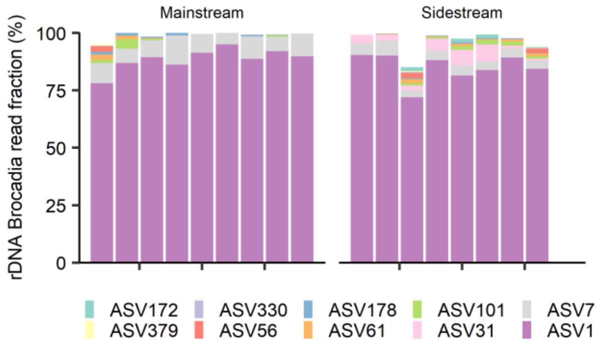


**Figure S5:** ASVs in the rDNA library classified as *Brocadia*. The top 10 AVS are shown.

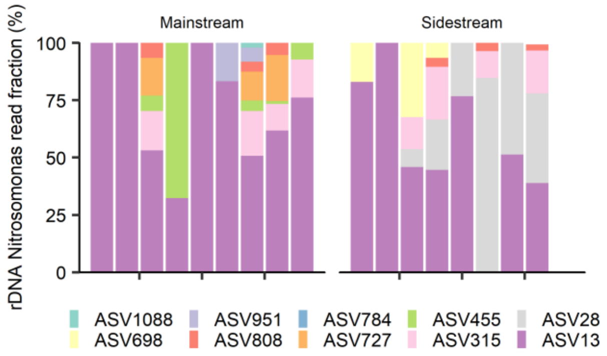


**Figure S6:** ASVs in the rDNA library classified as *Nitrosomonas*. The top 10 AVS are shown.

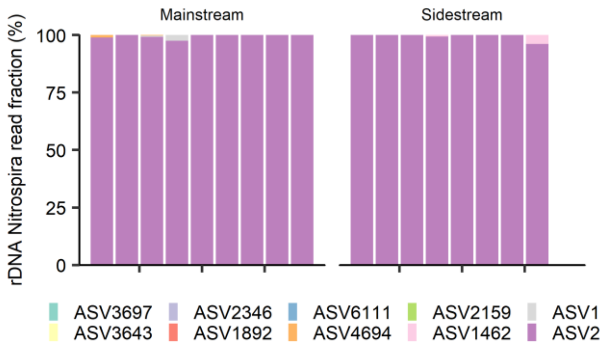


**Figure S7:** ASVs in the rDNA library classified as *Nitrospira*. The top 10 AVS are shown.

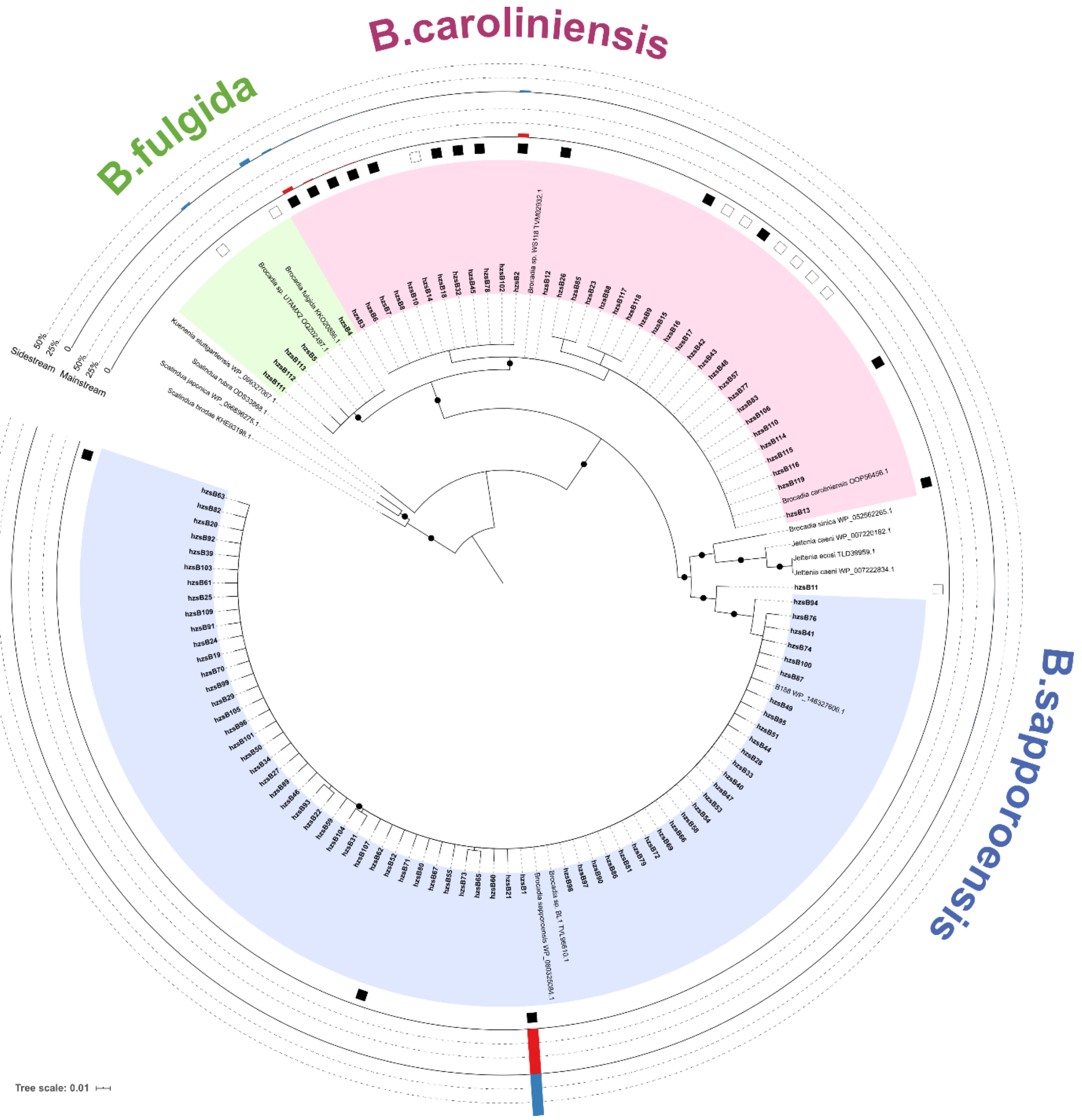

**Figure S8:** Anammox HzsB protein-based phylogenetic tree deduced from *hzsB* ASVs. Bootstrap values higher than 80% are indicated with a black circle. Bars indicate average relative abundance of the SV in mainstream (red) and sidestream (blue). Filled squares indicate ASVs with higher read abundance in mainstream, while empty squares indicate higher read abundance in sidestream (DESeq2; $p_{(adj)} < 0.01$).

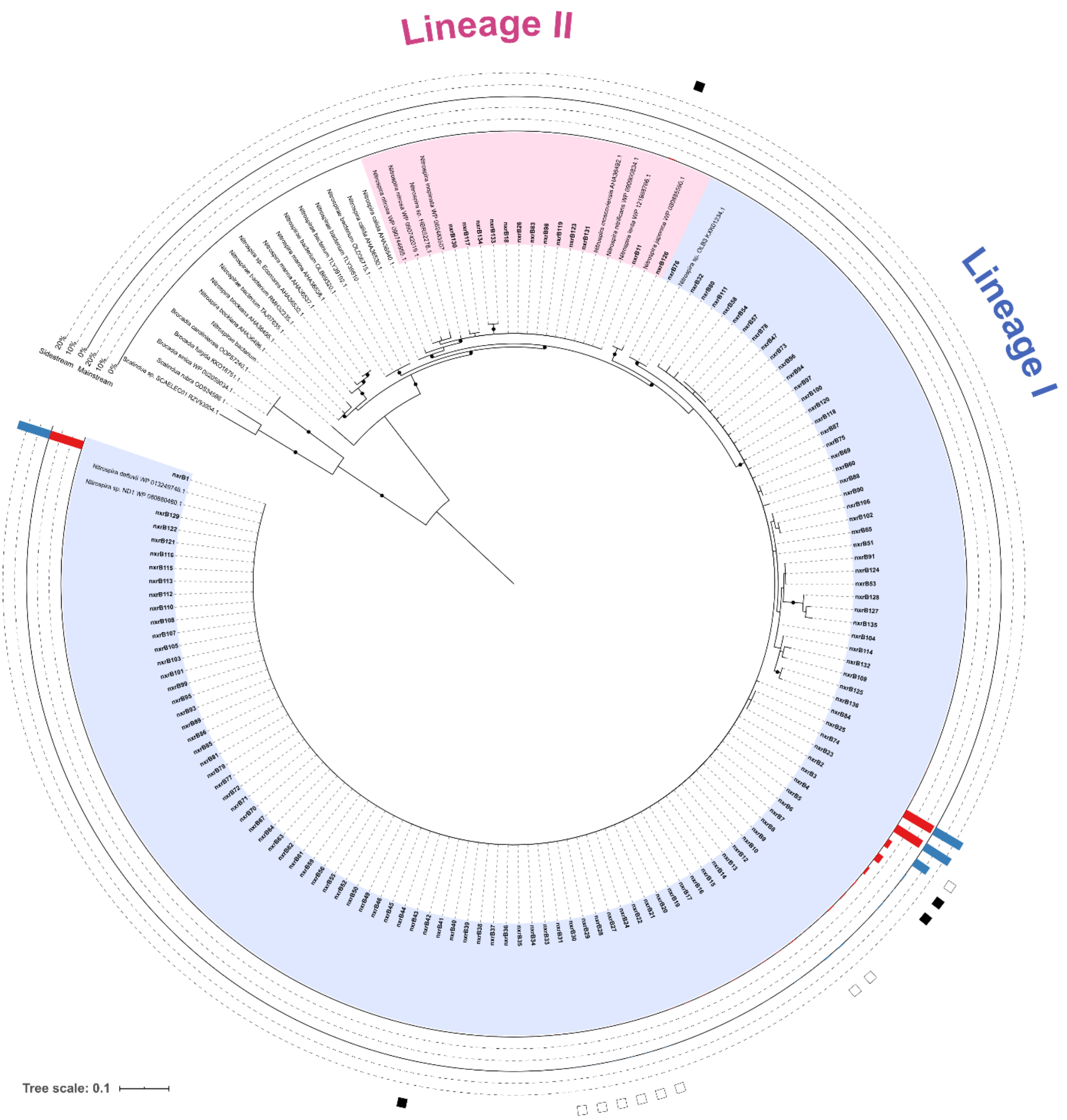


**Figure S9:** *Nitrospira* NxrB protein-based phylogenetic tree deduced from *NxrB* ASVs. Bootstrap values higher than 80% are indicated with a black circle. Bars indicate average relative abundance of the SV in mainstream (red) and sidestream (blue). Filled squares indicate ASVs with higher read abundance in mainstream, while empty squares indicate higher read abundance in sidestream (DESeq2; $p_{(adj)} < 0.01$). Lineage II was paraphyletic for *nxrB*.

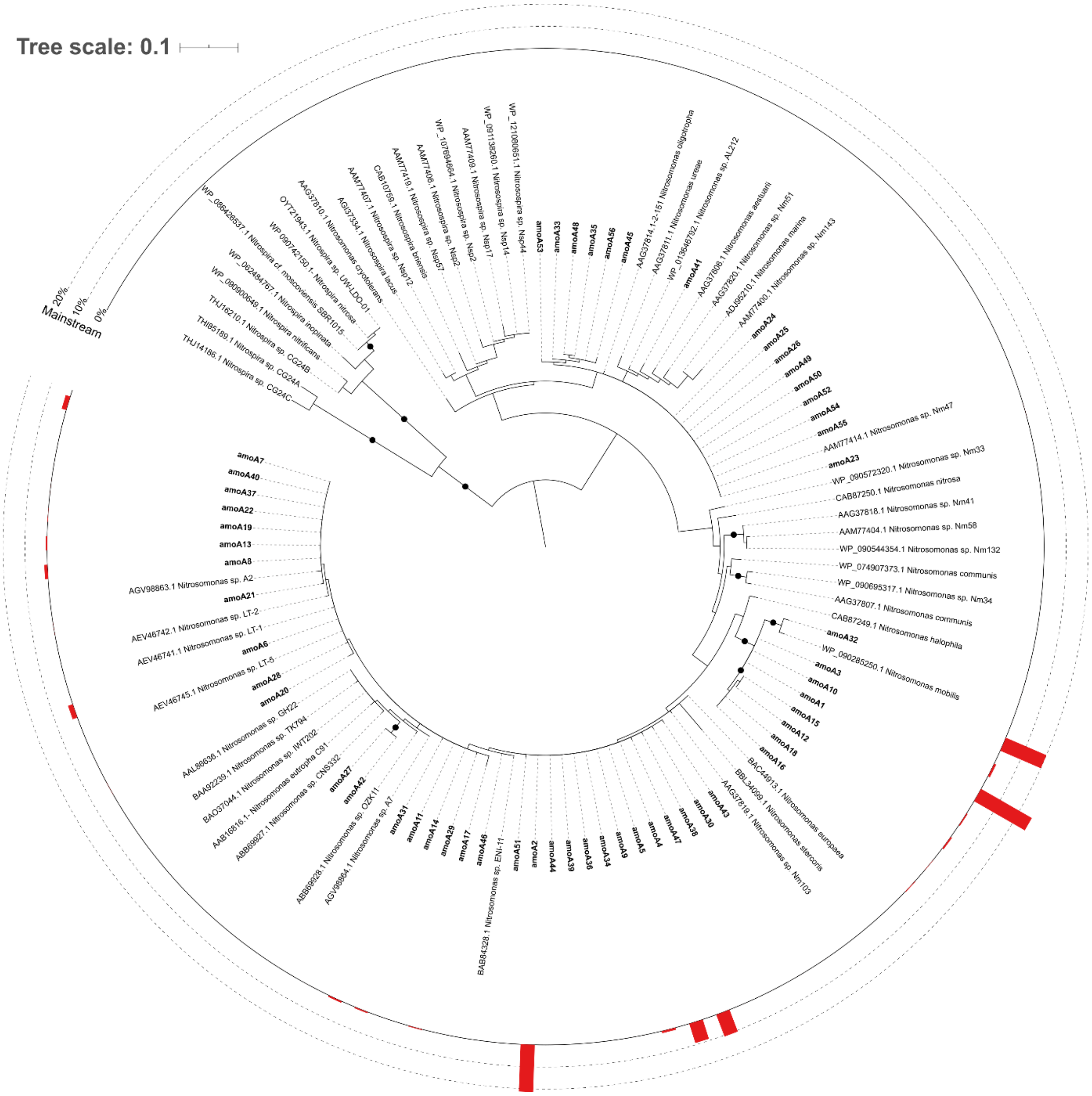


**Figure S10:** *Betaproteobacterial* AmoA protein-based phylogenetic tree deduced from *amoA* ASVs. Bootstrap values higher than 80% are indicated with a black circle. Red bars indicate average relative abundance of the ASV in mainstream.

**Supplementary tables**

**Table S1.** Accession list and metadata of Miseq reads. Carrier_ID indicates the biofilm carrier that was sampled. For example, for carrier M9 we did amplicon sequencing of 16S rDNA, 16s rRNA, *hzsB* DNA, *hzsB* mRNA, *nxrB* DNA and *amoA* DNA.

| Accession | Gene | Type | Reactor | Sampling_date | Sample_ID | Carrier_ID |
|---|---|---|---|---|---|---|
| SRS5064092 | 16S | DNA | Mainstream | 2015-10-15 | M_DNA_1 | M_1 |
| SRS5064087 | amoA | DNA | Mainstream | 2015-10-15 | M_DNA_1 | M_1 |
| SRS5064082 | hzsB | DNA | Mainstream | 2015-10-15 | M_DNA_1 | M_1 |
| SRS5064085 | 16S | RNA | Mainstream | 2015-10-15 | M_RNA_1 | M_1 |
| SRS5064092 | 16S | DNA | Mainstream | 2015-10-15 | M_DNA_10 | M_10 |
| SRS5064085 | 16S | RNA | Mainstream | 2015-10-15 | M_RNA_10 | M_10 |
| SRS5064092 | 16S | DNA | Mainstream | 2015-10-15 | M_DNA_11 | M_11 |
| SRS5064092 | 16S | DNA | Mainstream | 2015-10-15 | M_DNA_2 | M_2 |
| SRS5064091 | nxrB | DNA | Mainstream | 2015-10-15 | M_DNA_2 | M_2 |
| SRS5064082 | hzsB | DNA | Mainstream | 2015-10-15 | M_DNA_2 | M_2 |
| SRS5064087 | amoA | DNA | Mainstream | 2015-10-15 | M_DNA_2 | M_2 |
| SRS5064085 | 16S | RNA | Mainstream | 2015-10-15 | M_RNA_2 | M_2 |
| SRS5064092 | 16S | DNA | Mainstream | 2015-10-15 | M_DNA_3 | M_3 |
| SRS5064085 | 16S | RNA | Mainstream | 2015-10-15 | M_RNA_3 | M_3 |
| SRS5064092 | 16S | DNA | Mainstream | 2015-10-15 | M_DNA_4 | M_4 |
| SRS5064085 | 16S | RNA | Mainstream | 2015-10-15 | M_RNA_4 | M_4 |
| SRS5064092 | 16S | DNA | Mainstream | 2015-10-15 | M_DNA_5 | M_5 |
| SRS5064082 | hzsB | DNA | Mainstream | 2015-10-15 | M_DNA_5 | M_5 |
| SRS5064091 | nxrB | DNA | Mainstream | 2015-10-15 | M_DNA_5 | M_5 |
| SRS5064087 | amoA | DNA | Mainstream | 2015-10-15 | M_DNA_5 | M_5 |
| SRS5064091 | nxrB | DNA | Mainstream | 2015-10-15 | M_DNA_6 | M_6 |
| SRS5064087 | amoA | DNA | Mainstream | 2015-10-15 | M_DNA_6 | M_6 |
| SRS5064082 | hzsB | DNA | Mainstream | 2015-10-15 | M_DNA_6 | M_6 |
| SRS5064094 | hzsB | RNA | Mainstream | 2015-10-15 | M_RNA_6 | M_6 |
| SRS5064091 | nxrB | DNA | Mainstream | 2015-10-15 | M_DNA_7 | M_7 |
| SRS5064087 | amoA | DNA | Mainstream | 2015-10-15 | M_DNA_7 | M_7 |
| SRS5064082 | hzsB | DNA | Mainstream | 2015-10-15 | M_DNA_7 | M_7 |
| SRS5064085 | 16S | RNA | Mainstream | 2015-10-15 | M_RNA_7 | M_7 |
| SRS5064092 | 16S | DNA | Mainstream | 2015-10-15 | M_DNA_8 | M_8 |
| SRS5064091 | nxrB | DNA | Mainstream | 2015-10-15 | M_DNA_8 | M_8 |
| SRS5064087 | amoA | DNA | Mainstream | 2015-10-15 | M_DNA_8 | M_8 |
| SRS5064082 | hzsB | DNA | Mainstream | 2015-10-15 | M_DNA_8 | M_8 |
| SRS5064092 | 16S | DNA | Mainstream | 2015-10-15 | M_DNA_9 | M_9 |
| SRS5064091 | nxrB | DNA | Mainstream | 2015-10-15 | M_DNA_9 | M_9 |
| SRS5064087 | amoA | DNA | Mainstream | 2015-10-15 | M_DNA_9 | M_9 |
| SRS5064082 | hzsB | DNA | Mainstream | 2015-10-15 | M_DNA_9 | M_9 |
| SRS5064094 | hzsB | RNA | Mainstream | 2015-10-15 | M_RNA_9 | M_9 |
| SRS5064085 | 16S | RNA | Mainstream | 2015-10-15 | M_RNA_9 | M_9 |

| | | | | | | |
|---|---|---|---|---|---|---|
| SRS5064086 | 16S | DNA | Mainstream | 2015-06-09 | MO_DNA_1 | MO_1 |
| SRS5064086 | 16S | DNA | Mainstream | 2015-06-09 | MO_DNA_2 | MO_2 |
| SRS5064086 | 16S | DNA | Mainstream | 2015-06-09 | MO_DNA_3 | MO_3 |
| SRS5064086 | 16S | DNA | Mainstream | 2015-06-09 | MO_DNA_4 | MO_4 |
| SRS5064089 | 16S | DNA | Sidestream | 2015-10-15 | R_DNA_1 | R_1 |
| SRS5064088 | 16S | RNA | Sidestream | 2015-10-15 | R_RNA_1 | R_1 |
| SRS5064093 | nxrB | DNA | Sidestream | 2015-10-15 | R_DNA_2 | R_2 |
| SRS5064089 | 16S | DNA | Sidestream | 2015-10-15 | R_DNA_2 | R_2 |
| SRS5064088 | 16S | RNA | Sidestream | 2015-10-15 | R_RNA_2 | R_2 |
| SRS5064084 | hzsB | DNA | Sidestream | 2015-10-15 | R_DNA_3 | R_3 |
| SRS5064089 | 16S | DNA | Sidestream | 2015-10-15 | R_DNA_3 | R_3 |
| SRS5064088 | 16S | RNA | Sidestream | 2015-10-15 | R_RNA_3 | R_3 |
| SRS5064084 | hzsB | DNA | Sidestream | 2015-10-15 | R_DNA_4 | R_4 |
| SRS5064093 | nxrB | DNA | Sidestream | 2015-10-15 | R_DNA_4 | R_4 |
| SRS5064089 | 16S | DNA | Sidestream | 2015-10-15 | R_DNA_4 | R_4 |
| SRS5064088 | 16S | RNA | Sidestream | 2015-10-15 | R_RNA_4 | R_4 |
| SRS5064093 | nxrB | DNA | Sidestream | 2015-10-15 | R_DNA_5 | R_5 |
| SRS5064089 | 16S | DNA | Sidestream | 2015-10-15 | R_DNA_5 | R_5 |
| SRS5064084 | hzsB | DNA | Sidestream | 2015-10-15 | R_DNA_5 | R_5 |
| SRS5064090 | hzsB | RNA | Sidestream | 2015-10-15 | R_RNA_5 | R_5 |
| SRS5064088 | 16S | RNA | Sidestream | 2015-10-15 | R_RNA_5 | R_5 |
| SRS5064093 | nxrB | DNA | Sidestream | 2015-10-15 | R_DNA_6 | R_6 |
| SRS5064089 | 16S | DNA | Sidestream | 2015-10-15 | R_DNA_6 | R_6 |
| SRS5064084 | hzsB | DNA | Sidestream | 2015-10-15 | R_DNA_6 | R_6 |
| SRS5064088 | 16S | RNA | Sidestream | 2015-10-15 | R_RNA_6 | R_6 |
| SRS5064093 | nxrB | DNA | Sidestream | 2015-10-15 | R_DNA_7 | R_7 |
| SRS5064084 | hzsB | DNA | Sidestream | 2015-10-15 | R_DNA_7 | R_7 |
| SRS5064089 | 16S | DNA | Sidestream | 2015-10-15 | R_DNA_7 | R_7 |
| SRS5064088 | 16S | RNA | Sidestream | 2015-10-15 | R_RNA_7 | R_7 |
| SRS5064093 | nxrB | DNA | Sidestream | 2015-10-15 | R_DNA_8 | R_8 |
| SRS5064084 | hzsB | DNA | Sidestream | 2015-10-15 | R_DNA_8 | R_8 |
| SRS5064089 | 16S | DNA | Sidestream | 2015-10-15 | R_DNA_8 | R_8 |
| SRS5064090 | hzsB | RNA | Sidestream | 2015-10-15 | R_RNA_8 | R_8 |
| SRS5064083 | 16S | DNA | Sidestream | 2015-06-09 | RO_DNA_1 | RO_1 |
| SRS5064083 | 16S | DNA | Sidestream | 2015-06-09 | RO_DNA_2 | RO_2 |
| SRS5064083 | 16S | DNA | Sidestream | 2015-06-09 | RO_DNA_3 | RO_3 |
| SRS5064083 | 16S | DNA | Sidestream | 2015-06-09 | RO_DNA_4 | RO_4 |
| SRS5064083 | 16S | DNA | Sidestream | 2015-06-09 | RO_DNA_5 | RO_5 |
| SRS5064083 | 16S | DNA | Sidestream | 2015-06-09 | RO_DNA_6 | RO_6 |
| SRS5064083 | 16S | DNA | Sidestream | 2015-06-09 | RO_DNA_7 | RO_7 |
| SRS5064083 | 16S | DNA | Sidestream | 2015-06-09 | RO_DNA_8 | RO_8 |

**Table S2:** z-statistics and significance level for the beta regression model. See Figure S4, for plots.

| | z value | P-value | Significance |
|---|---|---|---|
| Intercept | -12.637 | < 2e-16 | *** |
| Log(DNA) | 17.978 | < 2e-16 | *** |
| Acidobacteria | -3.967 | 7.29E-05 | *** |
| Bacteroidetes | -0.923 | 0.356106 | |
| Chloroflexi | 2.532 | 0.011342 | * |
| Cyanobacteria | 1.579 | 0.114438 | |
| Firmicutes | 3.337 | 0.000847 | *** |
| Gemmatimonadetes | -1.004 | 0.315146 | |
| Nitrospirae | 1.11 | 0.266798 | |
| Planctomycetes | 10.155 | < 2e-16 | *** |
| Verrucomicrobia | -0.448 | 0.654134 | |
| Log(DNA)*Acidobacteria | -1.562 | 0.118306 | |
| Log(DNA)*Bacteroidetes | -0.068 | 0.946158 | |
| Log(DNA)*Chloroflexi | 2.258 | 0.023933 | * |
| Log(DNA)*Cyanobacteria | 2.501 | 0.012395 | * |
| Log(DNA)*Firmicutes | 1.911 | 0.056004 | . |
| Log(DNA)*Gemmatimonadetes | 0.178 | 0.858798 | |
| Log(DNA)*Nitrospirae | 1.157 | 0.247256 | |
| Log(DNA)*Planctomycetes | 8.671 | < 2e-16 | *** |
| Log(DNA)*Verrucomicrobia | 0.069 | 0.944744 | |

### References


Cribari-Neto F, Zeileis A. Beta Regression in R. *J STAT SOFTW* 2010; **34**: 24.

Ferrari S, Cribari-Neto F. Beta Regression for Modelling Rates and Proportions. *J Appl Statistics*. 2004; **31**: 799-815.

Letunic I, Bork P. Interactive Tree Of Life (iTOL) v4: recent updates and new developments. *Nucleic Acids Res*. 2019; **47**: W256–W259.

Pagès H, Aboyoun P, Gentleman R, DebRoy S. Biostrings: Efficient manipulation of biological strings. 2019

Schliep KP. phangorn: phylogenetic analysis in R. *Bioinformatics* 2010; **27**: 592-593.

Wright ES. Using DECIPHER v2.0 to Analyze Big Biological Sequence Data in R. *The R Journal*. 2016; **8**: 352-359.
